# Expression of the Close homolog of L-1 and embigin identifies activated myofibroblasts

**DOI:** 10.1101/2025.02.24.639984

**Authors:** Angus T Stock, Jess Day, Hannah Huckstep, Alexandra L Garnham, Ian P Wicks

**Affiliations:** WEHI, Melbourne, 3052, Australia.; University of Melbourne, Department of Medical Biology, 3052, Australia; Rheumatology Unit, The Royal Melbourne Hospital, 3050, Australia

## Abstract

Fibrosis is driven by the emergence of myofibroblasts, which are the primary producers of the extracellular matrix proteins that form fibrotic lesions. Despite this critical role, detecting myofibroblasts remains challenging due to the paucity of selective markers. We therefore screened for novel myofibroblast-specific markers, discovering that the expression of the close homolog of L1 (ChL-1) and embigin (Emb) distinguish activated myofibroblasts from their quiescent precursors. We report that ChL-1+/Emb+ fibroblasts: (1) emerge during cardiac inflammation, (2) have elevated expression of collagens and inflammatory factors and (3) localise to fibrotic zones - consistent with activated myofibroblasts. Mechanistically, Chl1+/Emb+ myofibroblasts differentiate from resident fibroblasts which upregulate these markers in response to proinflammatory cytokines, such as IL-1 and IL-17. Moreover, we show that embigin could be exploited to target antibody-based therapies to myofibroblasts and confirm this protein as a conserved marker of activated fibroblasts in multiple tissues and settings. Collectively, these findings identify ChL-1 and embigin as novel myofibroblast surface-markers that could be used to identify, enumerate and target this pathogenic population.

## Introduction

Fibrosis is defined as the excessive deposition of extracellular matrix (ECM) proteins (Henderson et al., 2020). It has been estimated that up to 45% of deaths in the United States of America are attributed to fibrotic disorders (Wynn, 2004). This alarming contribution to human mortality is attributed the fact that fibrosis: (1) develops in almost all serious and/or chronic diseases, and (2) directly impairs the homeostatic functions of all major organs (i.e. heart, lung, liver and kidney) by replacing parenchymal tissue and reducing tissue contractility (Distler et al., 2019, Henderson et al., 2020, Hinderer and Schenke-Layland, 2019). Consequently, fibrosis is now regarded as a most serious and common pathology.

Fibrosis is caused by the action of myofibroblasts, which are the primary cell type to produce and fold ECM proteins (Travers et al., 2016, Gibb et al., 2020, Ramachandran et al., 2019). While debate exists as to their origin (Hinz et al., 2007), it is now generally accepted that the myofibroblasts that drive interstitial fibrosis typically differentiate from resident, mesenchymal-derived fibroblasts that seed developing organs (Moore-Morris et al., 2014). Specifically, it is thought that local injury or inflammation trigger the production of pro-fibrotic factors such as TGFb, PDGF or interleukins (Henderson et al., 2020, Distler et al., 2019, Borthwick et al., 2013, Leask, 2010, Travers et al., 2016). These factors drive the activation of normally quiescent fibroblasts, which differentiate into activated myofibroblasts that produce the inflammatory and ECM proteins that execute tissue-remodelling (Henderson et al., 2020, Darby et al., 2014, Moore-Morris et al., 2014, Borthwick et al., 2013).

In the above model, the outcome of tissue-remodelling is determined by the magnitude and/or persistence of the myofibroblast response. Where injury or inflammation are mild and acute, myofibroblasts are short-lived and deposited ECM resolves to return to homeostasis (Darby et al., 2014, Hinz, 2007). However, in chronic or severe diseases, persisting inflammation leads to the sustained activation, expansion and/or survival of myofibroblasts. These events result in the excessive deposition of ECM, which form the unresolving fibrotic lesions that impair affected tissue (Darby et al., 2014, Henderson et al., 2020, Wynn, 2008).

Given the above pathogenesis, the ability to distinguish between quiescent fibroblasts and pathogenic myofibroblasts is vital. However, the capacity to discriminate between these population is remarkably limited, with the field largely relying upon alpha-smooth muscle actin (α-SMA) to identify myofibroblasts (Hinz et al., 2003, Hinz et al., 2007, Gibb et al., 2020). However, this protein has several limitations as a myofibroblast-specific marker. For instance, α-SMA is an intracellular protein (limiting live cell analysis/isolation), is expressed by other populations (such as pericytes and smooth muscle cells) and has been reported not to be expressed by myofibroblasts in tissues such as skeletal muscle (Zhao et al., 2018).

Given the above, we sought to identify novel cell surface proteins expressed by myofibroblasts. In doing so, we discovered that the transmembrane proteins Close homolog of L1 (ChL-1) and embigin (Emb) distinguish myofibroblasts from their quiescent precursors, revealing these proteins as novel myofibroblast-specific markers.

## Material & Methods

### Mouse models and injections

C57BL/6, BPSM-1 (Lacey et al., 2015), Wt1^CreERT2^ (Zhou et al., 2008) and R26^eYFP^ (Srinivas et al., 2001) were bred at the Walter and Eliza Hall Institute of Medical Research (WEHI, Australia) under specific pathogen-free conditions. The *Candida albicans* water-soluble (CAWS) complex was prepared as previously described (Nagi-Miura et al., 2006, Tada et al., 2008, Stock et al., 2016). To induce a Kawasaki-like disease, adult mice were injected intraperitoneally with 3mg CAWS on two consecutive days. For embryonic labelling of Wt1.eYFP mice, pregnant dams received tamoxifen (0.08mg/g dissolved in peanut oil) by oral gavage at E10.5. For in vivo labelling, mice were injected 10ug of anti-embigin PE (G7.43.1; ThermoFisher Scientific). To induce muscle injury, Barium Chloride (BaCl2) was dissolved in saline (at 1.2% weight/volume) and 50ul injected into the left quadricep of adult mice. All procedures were performed at WEHI and approved by the WEHI Animal Ethics Committee.

### Flow cytometric sorting, RNA-sequencing and bioinformatic analysis

For RNA-sequencing of cardiac fibroblasts, murine hearts were digested in type I collagenase (2mg/ml; Worthington) with DNase I (10μg/ml; Sigma) and single cell suspensions coated with anti-mouse Podoplanin-PE (8.1.1). Fibroblasts were then positively selected with the anti-PE microbeads on LS columns (Miltenyi), stained with directly conjugated mAbs against CD45.2 (104), CD31 (390), podoplanin (8.1.1) and PDGFRa/CD140A (APA5) and live fibroblasts (defined as propidium iodide negative single cells that were CD45-/CD31-/Podoplanin+/CD140A+) sorted on FACSAria II (BD) into RNAprotect Cell reagent (Qiagen). RNA was extracted using the RNeasy Micro Kit (Qiagen) and Illumina Stranded 150bp paired end mRNA sequencing performed by the Australian genome Facility (AGRF). For bioinformatic analysis, samples were aligned to the Mus musculus genome (mm39) using Rsubread (v2.10.2) (Lia et al., 2019). The proportion of fragments successfully mapped to the genome ranged from 97% to 99%. The number of fragments overlapping mouse genes was summarized using featureCounts with inbuilt annotation. Haemoglobin genes were removed, followed by the filtering of lowly expressed genes using edgeR’s filterByExpr function (Chen et al., 2016). The Xist gene was also removed in this step. After filtering, a total of 14,719 genes remained for downstream analysis. Differential expression (DE) analysis was performed using the limma (v3.46.0) (Ritchie et al., 2015) software package. Library sizes were normalized using the trimmed mean of M-values (TMM) method (Robinson & Oshlack., 2010). Differential expression was evaluated using limma-trend (Law et al., 2014) with robust empirical Bayes estimation of the variances (Phipson et al., 2016). The false discovery rate (FDR) was controlled below 0.05 using the method of Benjamini and Hochberg. Over-representation analysis of Gene Ontology (GO) terms and KEGG pathways for differentially expressed genes was conducted using limma’s goana and kegga functions. Mutli-dimensional scaling (MDS) plots and mean-difference (MD) plots were generated using limma’s plotMDS and plot MD functions respectively. Heatmaps were generated using pheatmap package.

### Flow cytometry of cardiac tissue

For flow cytometric analysis, murine hearts were digested as above and single cell suspensions stained with antibodies against CD45.2 (104), CD31 (390), podoplanin (8.1.1), PDGFRa/CD140A (APA5) and ChL-1 (polyclonal; AF2147), embigin (G7.43.1) and CSF2Rb (JORO50) (BD Bioscience, eBioscience, R&D systems or Biolegend). Cells were washed and stained with a secondary anti-goat 488 antibody to detect ChL-1 staining. Propidium iodide (100ng/ml) and cell counting beads were added immediately prior to data acquisition on a Fortessa (BD) FACS machine and analyzed using Flowjo software. For SAA3 analysis, hearts were digested in collagenase/DNAse containg Brefeldin A (5ug/ml), stained for surface markers as above, fixed, permeabilised (BD) and stained for intracellular SAA3 (JOR110A; BD) for analysis by flow cytometry.

### Real time quantitative PCR

For gene expression analysis, total fibroblasts or Chl1/Emb subsets were sorted for RNA extraction as above and cDNA synthesized with SuperScript III Reverse Transcriptase (Invitrogen) using oligo-dT primers (Promega). Quantitative real time PCR was performed with Fast Sybergren Master mix (In vitrogen) with customised primers described in Supplementary Table 1. Target gene expression was normalised to Hrpt (ΔCT), and relative expression converted by the 2-ΔCT method.

### Histology and immunofluorescent analysis

For imaging of mouse tissue, hearts were perfused with PBS and then fixed in 2% paraformaldehyde (4h on ice), dehydrated in 30% sucrose (12-18h at 4C) and embedded in OCT (Tissue-Tek). Hearts were sectioned (10μM) in the coronal plane and analyzed by histopathology or immunofluorescence microscopy. For histology, sections were stained with Sirius Red. For immunostaining, cardiac sections were hydrated in PBS, permeabilized with 0.1% Triton-X and non-specific staining blocked with serum (Jackson Immunoresearch), 1% BSA and Protein block (Dako). Sections were stained with primary antibodies against GFP (ab290; Abcam), ChL-1 (polyclonal; AF2147) and embigin (G7.43.1) before detection with fluorochrome conjugated secondary antibodies (Invitrogen or Abcam). Slides were counterstained with DAPI, imaged on a Zeiss LSM-880 Confocal Microscope and analyzed with ImageJ software.

### In vitro assays

To generate primary cardiac fibroblasts, hearts were digested in type I collagenase/DNase I and cardiac cells cultured in DMEM containing 10% foetal bovine serum. When cells had reached 30% confluence, fibroblasts were treated with one of IL-1b, IL-17, Tnf, Tgfb, IFNg, GM-CSF, FGF or PDGFbb (20ng/ml; Peprotech) daily for two days. Cells were then harvested with trypsin and stained for embigin and podoplanin for analysis by flow cytometry as above.

## Results

### The Close homolog of L1 and Embigin are upregulated by a subset of cardiac fibroblasts during fibrosis

To identify novel myofibroblast-specific markers, we first screened for transmembrane proteins upregulated by fibroblasts during fibrosis. To do so, we utilised two forms of cardiac fibrosis. The first is a mouse model of Kawasaki Disease (KD), where cardiac vasculitis is induced by the injection of the yeast wall fraction of Candida albicans, referred to as Candida albicans water soluble complex (CAWS). The second is the Bone Phenotype Spontaneous Mutant (BPSM) mouse line, which have a retrotransposon insertion in the 3’ untranslated region of *Tnf*, resulting in the constitutive hyperproduction of TNFa (Lacey et al., 2015). Histological analysis confirmed that both CAWS injected mice (∼4 weeks post injection) and BPSM heterozygous mice (∼10-14 weeks of age) develop extensive interstitial cardiac fibrosis localising to the aortic root and epicardial adipose tissue surrounding the coronary arteries (**Fig. 1A**). Hence, these systems provide useful models to study cardiac fibrosis and the myofibroblasts that drive this pathology.

**Figure 1.**
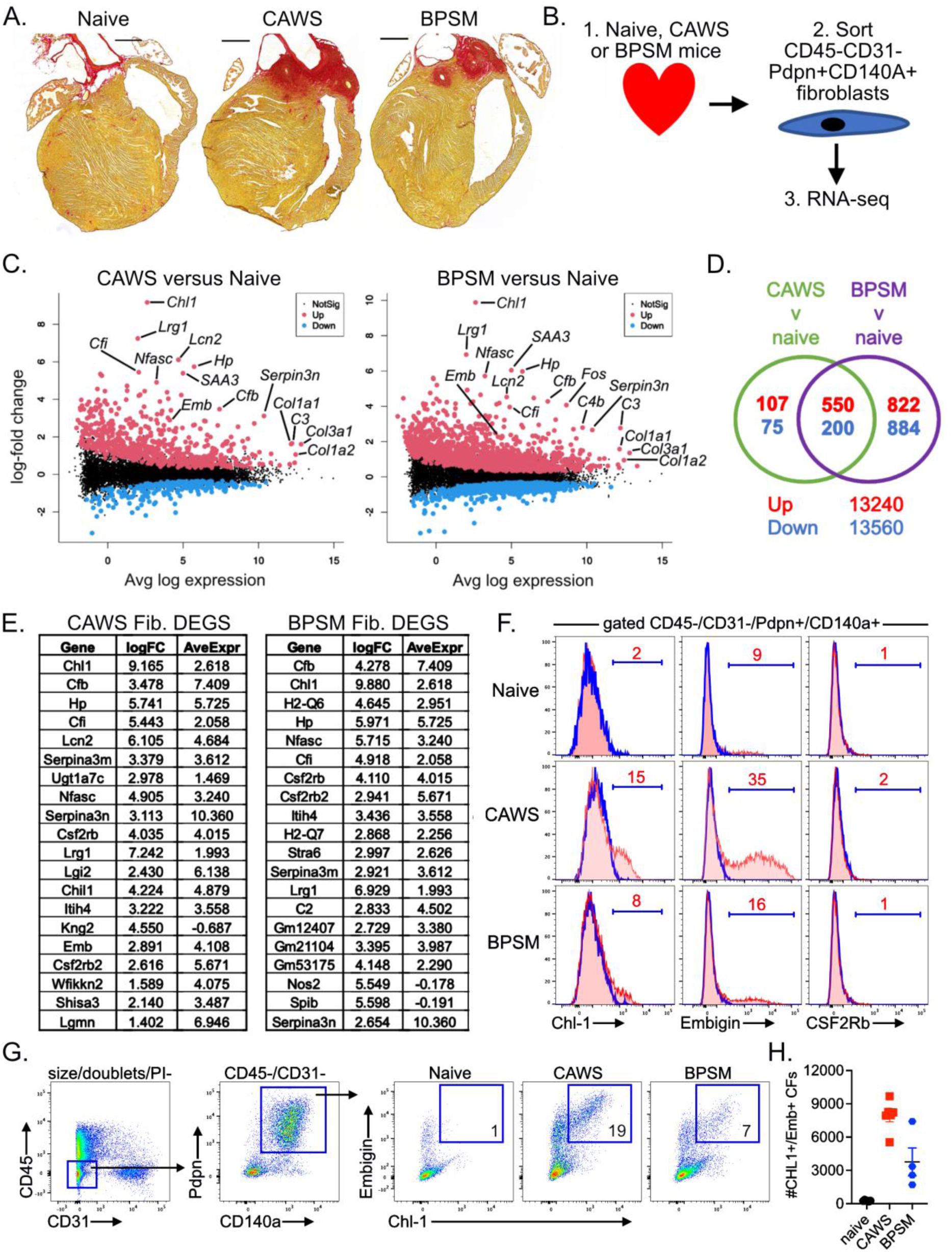
ChL-1+/Emb+ fibroblasts emerge during cardiac fibrosis. **(A)** Sirius red stained cardiac sections from naïve or CAWS injected wild-type mice or BPSM-1 mice. **(B)** Experimental schema for RNA-sequencing. **(C)** Volcano plots with selected conserved differentially expressed genes (DEGs) annotated. **(D)** Venn diagrams show the number of common and unique DEGS in each group. **(E)** Twenty top-ranked DEGS for each group. **(F)** Histograms gated on cardiac fibroblasts (CD45-CD31-CD140A+Pdpn+) from naïve, CAWS and BPSM mice show the expression of RNA-seq identified surface proteins ChL-1, embigin and CSFR2b. The blue line shows control staining (FMO or secondary alone) and red line shows test antibody, with the percentage of fibroblasts positive for each marker inset. **(G-H)** Flow cytometry of hearts show gating strategy and enumeration ChL-1/Emb fibroblasts for naïve, CAWS and BPSM mice. Dots depict individual mice pooled from two independent experiments.

To identify genes upregulated during fibrosis, we performed RNA-sequencing (RNA-seq) on fibroblasts (characterised by CD45-/CD31-/Podoplanin+/CD140a+ expression) sorted from the hearts of naïve, CAWS injected (∼4 weeks post injection) and BPSM (at ∼10-14 weeks of age) mice (**Fig. 1B**). Bioinformatic analysis then identified genes that were differentially expressed in CAWS or BPSM cardiac fibroblasts relative to their quiescent, naïve counterparts. This analysis revealed that fibrosis was associated with a profound transcriptional change, with fibroblasts from CAWS injected mice increasing the expression of ∼650 genes relative to their naïve counterparts and BPSM fibroblasts increasing ∼1350 genes (**Fig. 1C-D**). Notably, a vast majority of upregulated, differentially expressed genes (DEGs) identified in CAWS fibroblasts were also upregulated in BPSM mice (550 of 657 genes), suggesting that fibroblasts enter a relatively conserved transcriptional program in both disease settings (**Fig. 1C-D)**.

Our RNA-seq data revealed a number DEGSs encoding transmembrane proteins which speculated may serve as myofibroblast-specific surface markers. These include the cell adhesion molecule *the Close-homolog of L1* (ChL-1), which was the top-ranked DEG in the CAWS group and ranked second in the BPSM group (**Fig. 1E**). Notably, a second member of the L1 family, neurofascin (Nfasc), was also highly ranked in both CAWS (8^th^ ranked DEG) and BPSM (5^th^ ranked DEG) groups, indicating that the L1 molecules may play a prominent role in myofibroblast biology (Wei and Ryu, 2012). Other surface proteins were also highly upregulated in both fibrotic groups, including the IgG superfamily member embigin (Emb) and the beta chain of the CSF2 receptor (CSF2Rb/CD131).

We next validated the expression of candidate genes by flow cytometry, staining cardiac fibroblasts from naïve, CAWS or BPSM mice for ChL-1, Embigin and CSF2-Rb. While CSF2Rb was not detected by any fibroblasts tested, this analysis revealed that both ChL-1 and embigin was substantially increased on fibroblasts from CAWS and BPSM mice compared to naïve controls (**Fig. 1F**). Dual expression analysis confirmed that ChL-1 and embigin were co-expressed, with a distinct population of ChL-1+/Emb+ fibroblasts emerging within the hearts of CAWS injected and BPSM mice (**Fig. 1G-H**). Collectively, this analysis reveals that the surface proteins Close homolog of L1 (ChL-1) and Embigin are upregulated by a subset of cardiac fibroblasts that emerge during fibrosis.

### ChL-1+/Emb+ fibroblasts express fibrotic and inflammatory factors and co-localise with fibrosis

We next investigated whether ChL-1+/Emb+ fibroblast subset were myofibroblasts. A defining feature of myofibroblasts is their elevated production of extracellular matrix (ECM) proteins and inflammatory factors (Gibb et al., 2020, Distler et al., 2019, Henderson et al., 2020). Consistent with this concept, our RNA-seq analysis revealed that genes encoding collagens, collagen-folding enzymes (such as P4HA3) and inflammatory factors (such as complement factors, cytokine and chemokines) were significantly increased by fibroblasts from fibrotic CAWS and BPSM mice (**Fig. 1**). We therefore determined if these factors were predominantly expressed by ChL-1+/Emb+ fibroblasts. To this end, we sorted either total cardiac fibroblasts from naïve or CAWS injected mice or further divided fibroblasts (from CAWS mice) into ChL-1-/Emb- and ChL-1+/Emb+ subsets before measuring candidate genes by RT-PCR (**Fig. 2A**). This analysis revealed that ChL-1+/Emb+ fibroblasts showed highly elevated expression of ECM and inflammatory factors compared to either ChL-1-/Emb- or total fibroblasts. Specifically, ChL-1+/Emb+ fibroblasts isolated from CAWS mice expressed ∼3-5 fold higher levels of the fibrillar collagens 1a1/1a2/3, ∼10-fold higher expression of P4Ha3 (an enzyme responsible for collagen folding) and substantially higher levels of SAA3, haptoglobin, complement factors and chemokines. Moreover, ex vivo intracellular staining (ICS) for SAA3 confirmed (at a protein level) that fibroblasts produce the inflammatory cytokine SAA3 upon CAWS induced cardiac disease (**Fig. 2D**), and more critically, SAA3 production was restricted to ChL-1+/Emb+ fibroblasts. Thus, ChL-1+/Emb+ expression identifies fibroblasts with an activated transcriptional program, characterised by the elevated expression collagens, collagen-folding enzymes and inflammatory factors.

**Figure 2.**
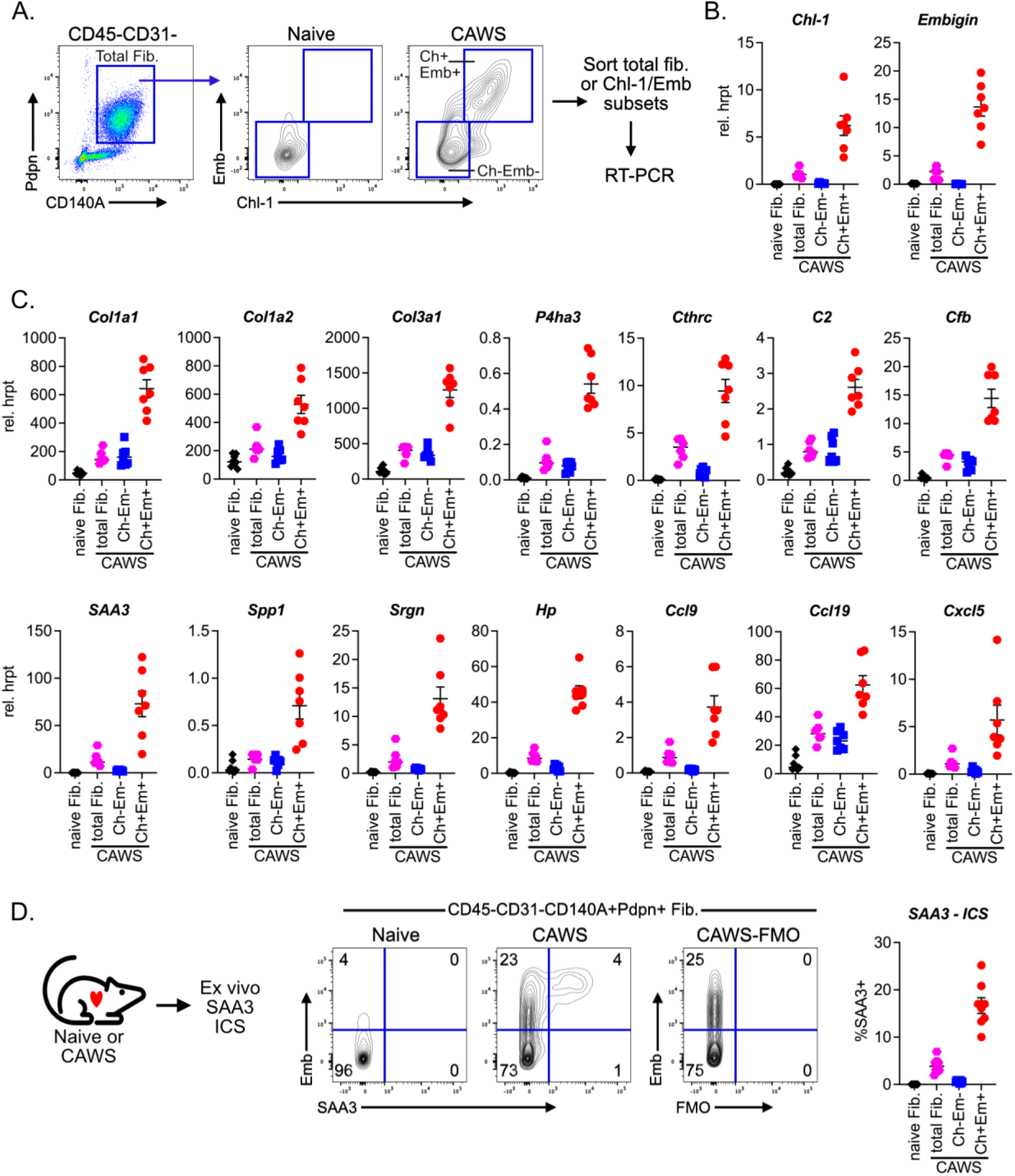
ChL-1+/Emb+ fibroblasts express fibrotic and inflammatory factors. **(A-C)** Total fibroblasts (CD45-CD31-CD140A+Pdpn+) or ChL-1+/Emb+ and ChL-1-/Emb-subsets were sorted from the hearts of naïve or CAWS injected mice for analysis by RT-PCR. **(A)** Flow cytometric plots show gating strategy for sorting and graphs depict expression of marker genes **(B)** and fibrotic and inflammatory factors **(C)**. Dots depict values from individual or pools of mice (2-3/group) acquired in two independent experiments. **(D)** Hearts from naïve or CAWS injected mice (3-4 weeks post injection) were analysed by intracellular staining (ICS) for SAA3. Contour plots are gated on CD45-CD31-CD140A+Pdpn+ cardiac fibroblasts while graphs depict %SAA3+ for total fibroblasts or ChL-1/Emb subsets. Dots depict individual mice pooled from two independent experiments.

We next determined if ChL-1+/Emb+ fibroblasts localised to fibrosis. To explore this question, we made use of the fact that CAWS-induced fibrosis is largely restricted to the aortic root and upper regions of the heart (**Fig. 3A**). We therefore dissected the hearts into the upper and lower regions to obtain fibrotic and non-fibrotic tissue respectively. Flow cytometric analysis of defined regions revealed that ChL-1+/Emb+ fibroblasts were abundant within the upper region of the heart where fibrosis develops and entirely absent from non-fibrotic lower tissue (**Fig. 3B**). Collectively, the above findings illustrate that ChL-1 and Emb is expressed by a subset of fibroblasts that: (1) emerge during cardiac disease, (2) express ECM and inflammatory factors and (3) localise to fibrosis. Consequently, we propose that ChL-1+/Emb+ expression identifies the activated myofibroblasts that drive fibrosis

**Figure 3.**
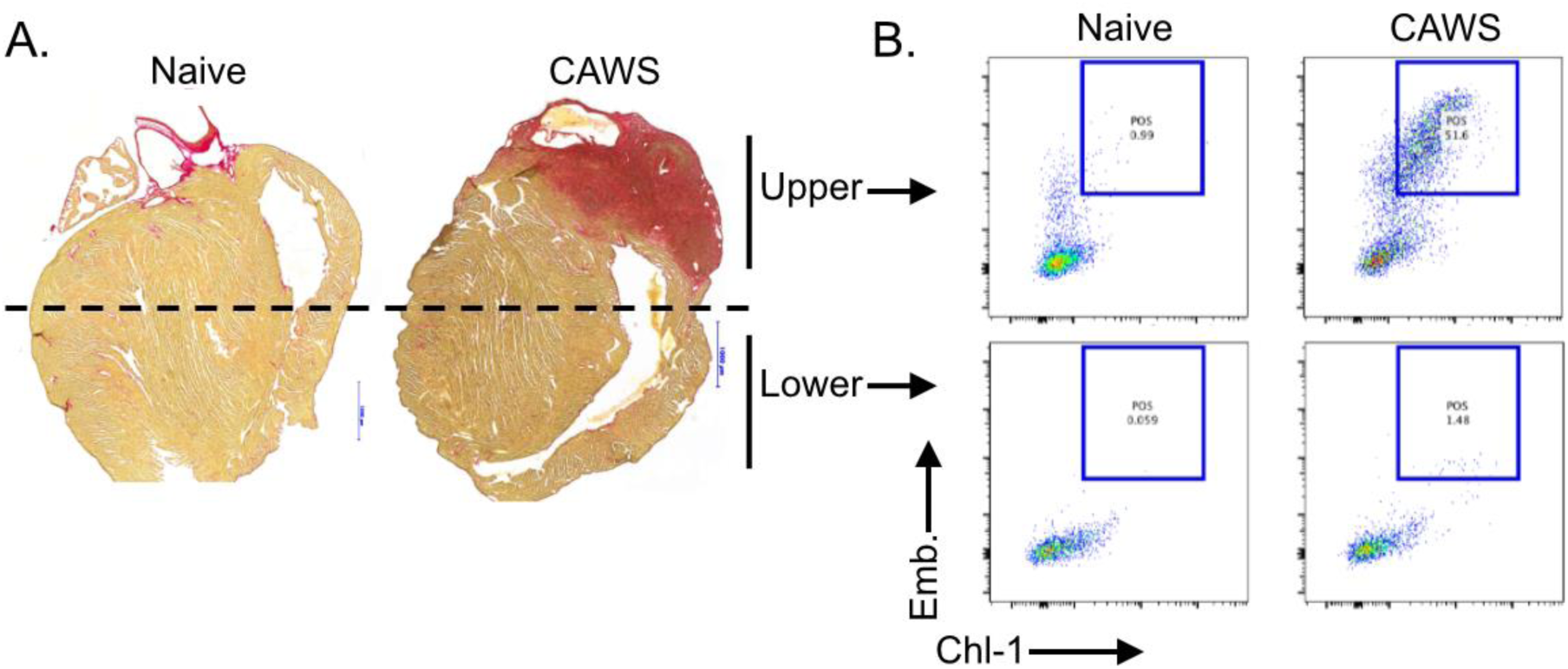
ChL-1+/Emb+ fibroblasts localise to fibrotic zones within the heart. **(A)** Sirius red stained cardiac sections from naïve and CAWS injected (4-5 weeks post injection) mice. **(B)** Flow cytometry of cardiac tissue from upper or lower cardiac tissue showing enumeration of ChL-1/Emb fibroblasts from naïve and CAWS mice. Representative image is shown of 2-3 independent experiments.

### Epicardial-derived, resident fibroblasts upregulate ChL-1 and Embigin during cardiac disease

We next investigated the origin of ChL-1+/Emb+ myofibroblasts, asking if they develop from resident fibroblasts. To do so, we performed lineage tracing with the Wilms Tumor 1 (Wt1) CreERT2 mouse line. Wt1 is a transcription factor that is temporally restricted to the epicardium during embryogenesis, which are the major source of cardiac fibroblasts (von Gise et al., 2011). Consequently, embryonic WT1-driven recombination with Wt1^CreERT2^ mice has been widely used to achieve the indelible labelling of epicardial-derived, resident cardiac fibroblasts (Zhou and Pu, 2012, Moore-Morris et al., 2014). We used this approach to determine if the resident fibroblasts the seed the heart during development upregulate ChL-1 and Emb during cardiac inflammation. To this end, we crossed Wt1^CreERT2^ and R26^floxed-stop-eYFP^ reporter mice to produce Wt1^CreERT2^.R26^eYFP^ mice (**Fig. 4A**). In this system, tamoxifen administration at E10.5 of gestation results in eYFP labelling of the Wt1+ epicardium, which in turn, give rise to eYFP+ resident cardiac fibroblasts (Zhou and Pu, 2012).

**Figure 4.**
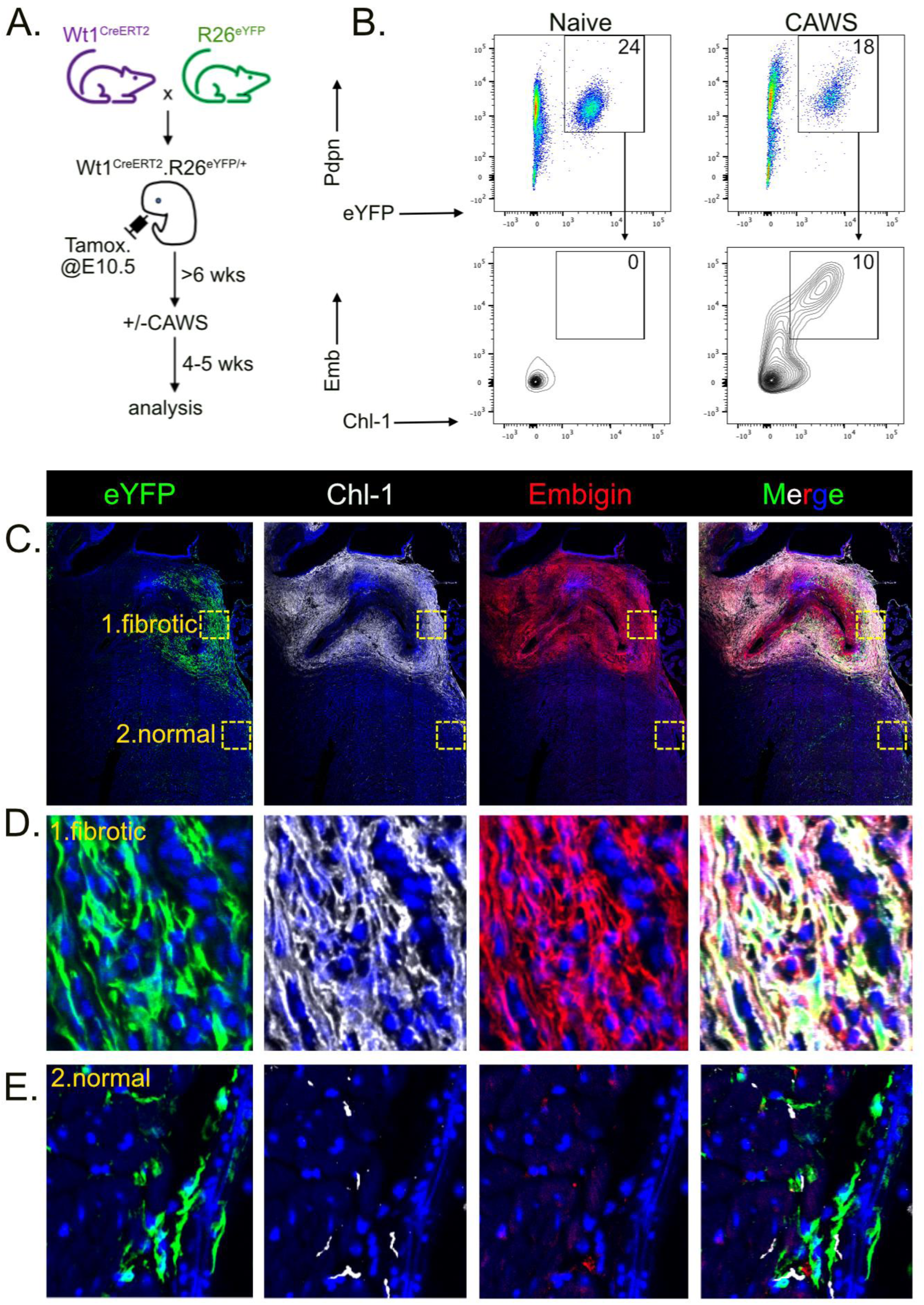
Epicardial-derived fibroblasts upregulate ChL-1 and embigin during cardiac fibrosis. **(A)** Experimental schema. **(B)** Hearts from naïve and CAWS injected (4-5 weeks post injection) Wt1^CreERT2^.R26^eYFP^ mice analysed by flow cytometry. Pseudeo-density plots are gated on CD45-CD31-events and contour plots show ChL-1 and embigin expression by CD45-CD31-Pdpn+eYFP+ epicardial-derived fibroblasts. **(C-E)** Cardiac sections from a CAWS injected Wt1^CreERT2^.R26^eYFP^ mice stained for eYFP (green), ChL-1 (white), Embigin (red) and DAPI (blue). Inset area show fibrotic regions **(D)** or normal tissue **(E)**. Representative data is shown of 2-3 independent experiments.

For our experiments, tamoxifen treated Wt1^CreERT2^.R26^eYFP^ mice were challenged with CAWS and 4-5 weeks later (when substantial cardiac fibrosis has developed) analysed by flow cytometry. The analysis of hearts from naïve mice revealed that in the steady-state, eYFP+ epicardial-derived, resident cardiac fibroblasts did not express ChL-1 or Emb. However, upon CAWS challenge, a subset of eYFP+ resident cardiac fibroblasts clearly upregulated ChL-1 and Emb (**Fig. 4B**). We next determined if these ChL-1+/Emb+ resident fibroblasts localise to fibrotic zones by analysing cardiac sections by confocal microscopy. This analysis revealed that eYFP+ fibroblast had expanded in the fibrotic areas of the heart, and critically, stained positive for both ChL-1 and embigin (**Fig. 4C**). In contrast, the eYFP+ fibroblast that populated distal areas of normal cardiac tissue did not express these markers. Collectively, these findings show that the ChL-1+/Emb+ myofibroblasts that emerge and localise with fibrosis differentiate from ChL-1-/Emb-epicardial-derived, resident fibroblasts that seed the heart during gestation.

### Inflammatory cytokines induce Embigin expression on cardiac fibroblasts *in vitro*

Our findings show that ChL-1 and embigin are expressed as fibroblasts differentiate from a quiescent to myofibroblast state. We next examined potential regulators of ChL-1 and embigin expression, exploring the possibility that these are cytokine inducible proteins. To test this, we treated primary cultured cardiac fibroblasts with various cytokines for 48 hours before measuring embigin expression. We found that while the putative pro-fibrotic cytokine Tgfb had little effect, the pro-inflammatory cytokines IL-1b and IL-17 robustly induced embigin expression on fibroblasts (**Fig. 5A-B**). Notably, we did not detect ChL-1 expression, although speculate that this protein may have been cleaved off by trypsin when harvesting fibroblasts (data not shown). These findings raise the possibility that Embigin expression identifies myofibroblasts that are activated by the pro-inflammatory cytokines, such as IL-1b and IL-17, that emerge during cardiac disease.

**Figure 5.**
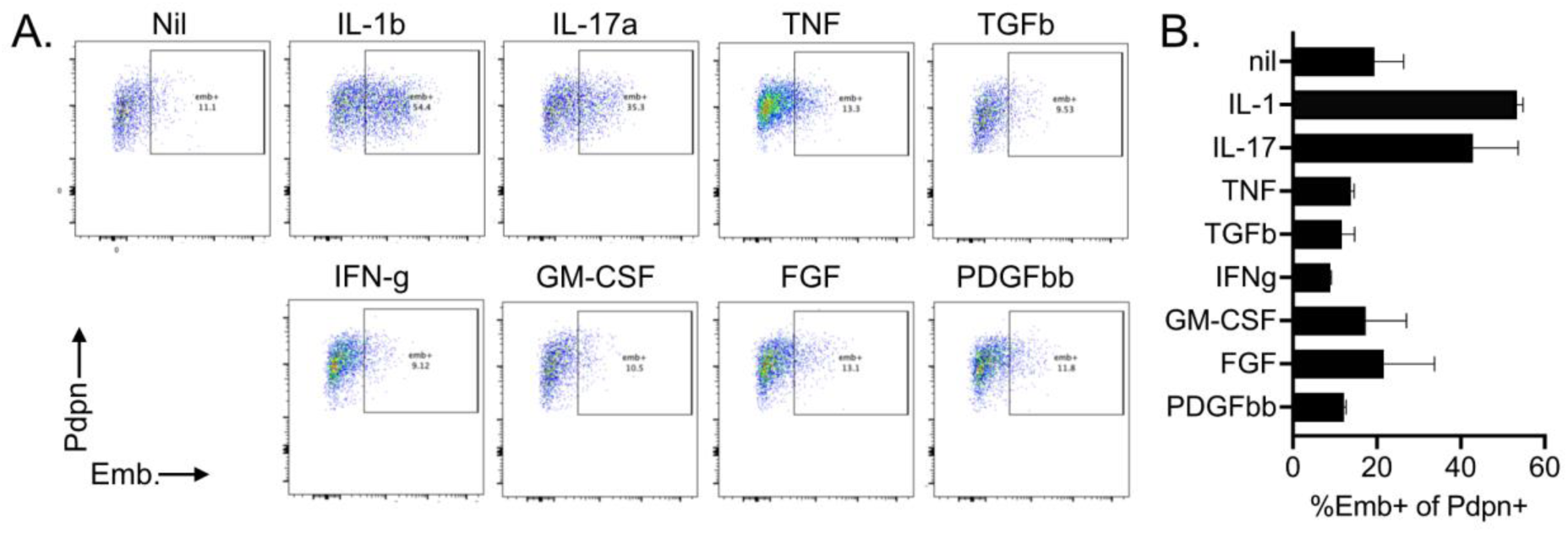
Pro-inflammatory cytokines induce embigin expression on fibroblasts. **(A)** Primary cardiac fibroblasts were established in culture and treated with various cytokines for 48 hours before the expression of embigin was examined on podoplanin+ fibroblasts. **(B)** Graphs show mean±SEM pooled data from two independent.

### Parenterally administered anti-Embigin antibodies bind myofibroblasts *in vivo*

The above findings illustrate that ChL-1 and embigin are expressed by myofibroblasts. One potential application for these findings is that antibodies against these proteins could be used to target therapies to pathogenic myofibroblasts. To evaluate this potential, we examined if anti-embigin antibodies could bind cardiac myofibroblasts when injected into fibrotic mice? To this end, we used the CAWS injection to induce cardiac fibrosis before administering a fluorochrome (PE) conjugated anti-Emb antibody. Specifically, ant-Emb antibody was injected intravenously into fibrotic mice and 30 minutes later, hearts were examined by flow cytometry (**Fig. 6A**). By using ChL-1 expression to distinguish quiescent fibroblasts (ChL-1-) and activated myofibroblasts (ChL-1+), we found that while quiescent fibroblasts showed minimal labelling, ChL-1+ myofibroblasts were clearly bound by the intravenously injected anti-Emb antibody (**Fig. 6B**). These findings reveal that anti-embigin antibodies can access the inflamed heart and selectively coat myofibroblasts.

**Figure 6.**
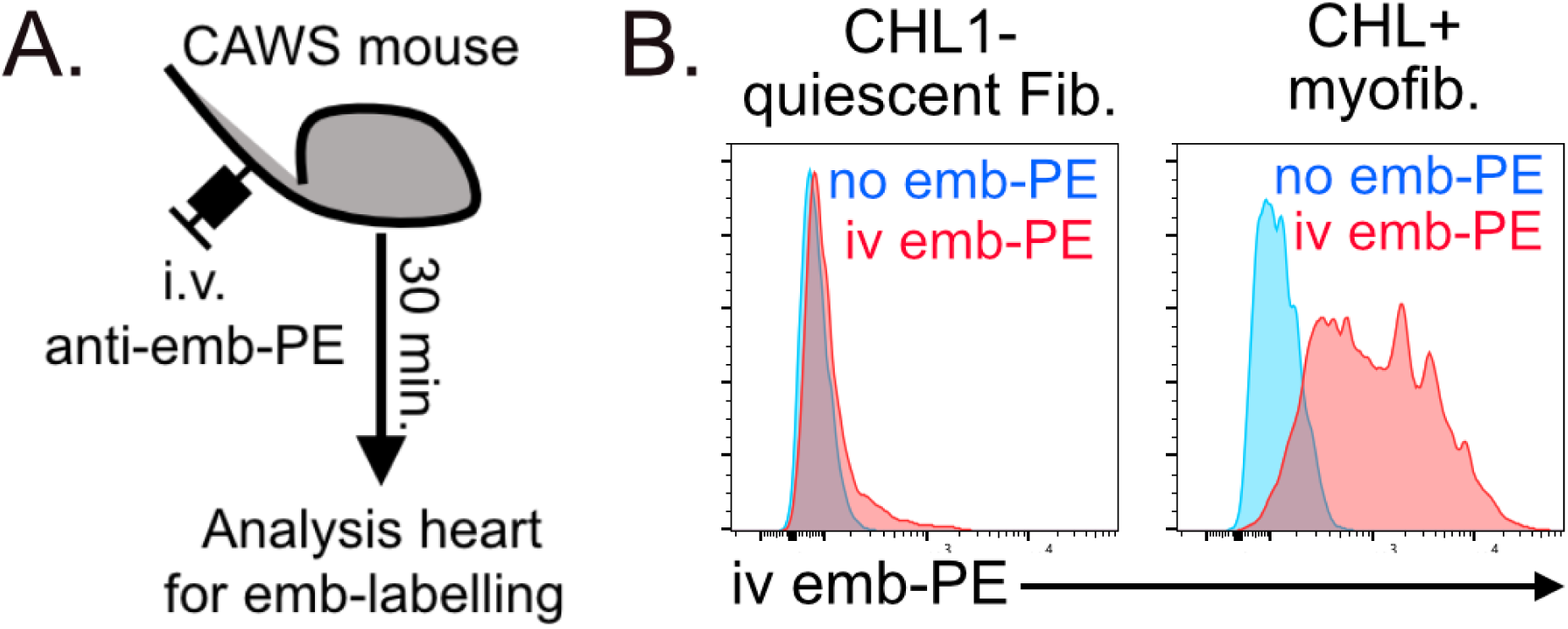
Parenterally administered anti-embigin antibodies can bind myofibroblasts in vivo. **(A)** Experimental schema. **(B)** Histogram plots show labelling by intravenously injected anti-embigin PE antibody on ChL-1- and ChL-1+ cardiac fibroblasts from CAWS mice. A representative plot from two independent experiments (n=5) is shown.

### Fibroblasts upregulate embigin during acute skeletal muscle injury

The above shows that ChL-1 and embigin expression identify the myofibroblasts that drive cardiac fibrosis in heart diseases. These findings raise the question of whether activated fibroblasts in non-cardiac tissue, and/or other disease settings, also express these markers? To address this question, we utilised the barium chloride (BaCl2)-induced muscle injury model, a popular method for inducing muscle necrosis and inflammation within skeletal muscle (Jung et al., 2019). Specifically, we injected BaCl2 into the quadricep of mice and at various time-points, analysed the expression of ChL-1 and embigin on muscle fibroblasts. This time-course revealed that fibroblasts expand and increase Podoplanin expression following BaCl2 injury, before returning to baseline levels at day 17 (**Fig. 7**). Critically, we found that fibroblast expansion coincided with the upregulation of embigin, which peaked at 4 to 7 days post injury, before largely returning to baseline levels at day 17 (**Fig. 7**). Intriguingly, despite embigin upregulation, we observed relatively minimal expression of ChL-1, with most fibroblasts exhibiting an intermediate ChL-1-/Emb+ phenotype. Despite the differential ChL-1 expression, these findings clearly show that embigin is transiently expressed on by fibroblasts in injured skeletal muscle, suggesting that this marker may be broadly useful in identifying acutely activated fibroblasts in both cardiac and skeletal muscle and possibly, other tissues.

**Figure 7.**
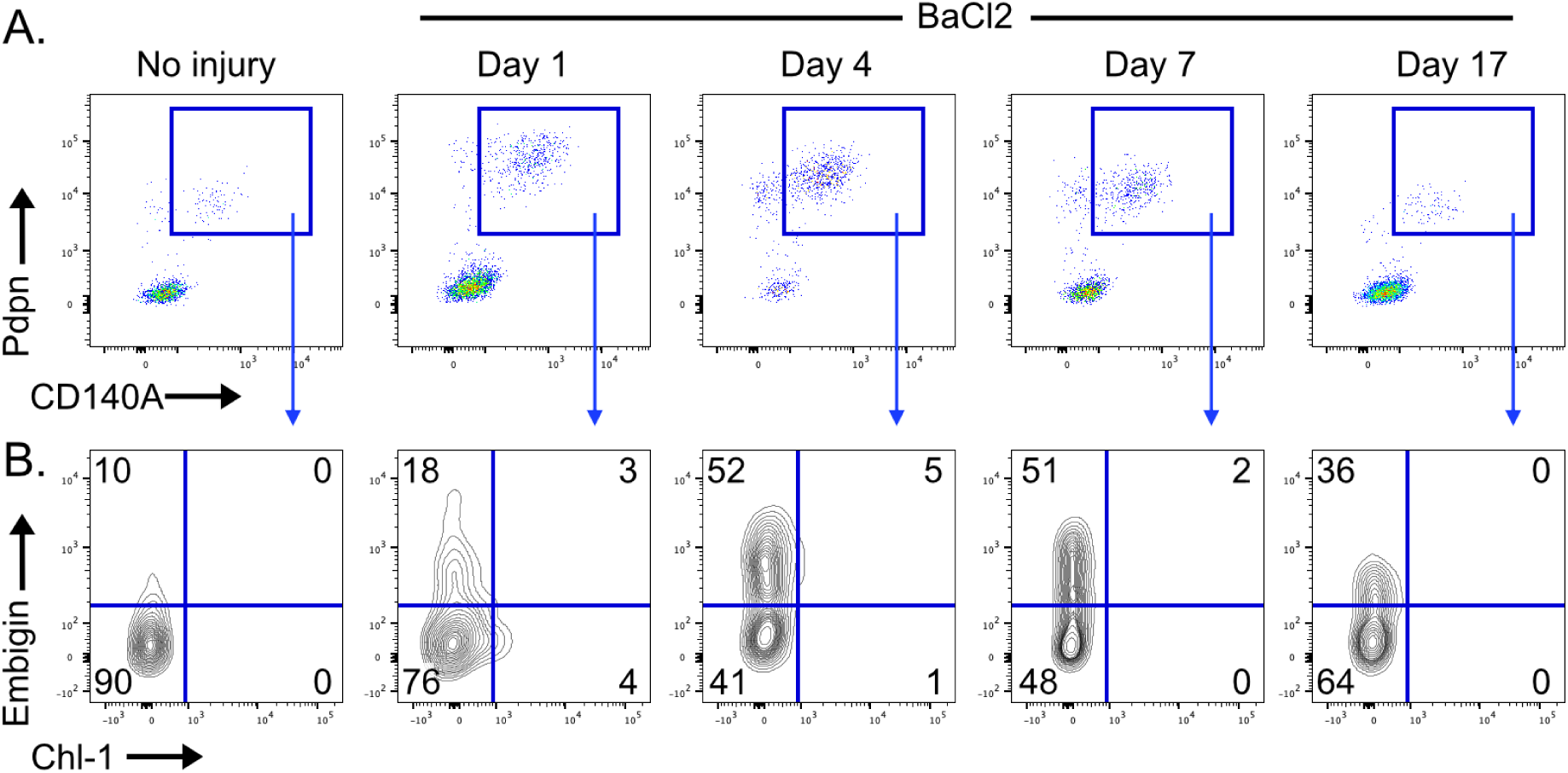
Muscle fibroblasts upregulate embigin during acute injury. **(A-B)** BaCl2 was injected into the left quadricep of mice before analysis by flow cytometry. **(A)** Dot-plots are gated on llive CD45-/CD31-events and **(B)** contour plots show ChL-1 and embigin expression by muscle fibroblasts. A representative plot from at least two independent experiments (n>5 per time-point) is shown.

## Discussion

Here, we have identified two novel surface markers that distinguish activated myofibroblasts from their quiescent precursors. Specifically, we report that the co-expression of the Close homolog of L1 (Chl1) and embigin (Emb) identify fibroblasts that: (1) emerge during cardiac inflammation, (2) have elevated expression of collagens and inflammatory factors and (3) localise to fibrotic zones. Given these features, we propose that ChL-1 and Emb expression identifies the activated myofibroblasts that drive fibrosis.

Our findings raise the question of whether ChL-1 and embigin control myofibroblast function? While these are sparsely studied proteins, it is intriguing to note that both ChL-1 and embigin are cell adhesion molecules most widely reported to regulate neuronal cell behaviour such as migration, differentiation and sprouting (Guseva et al., 2018, Lain et al., 2009). In addition, we found that NFASC - another member of the L1 family (Wei and Ryu, 2012) - is also upregulated on activated fibroblasts. Thus, at face value it appears that myofibroblasts express multiple neuronal mobility proteins. However, why this is the case and whether these proteins control myofibroblasts biology remains to determined.

We have used lineage tracing to investigate the origin of ChL-1+/Emb+ myofibroblasts. Our findings reveal that epicardial-derived, resident cardiac fibroblasts that seed the heart during embryogenesis upregulate ChL-1 and Emb expression during cardiac disease. Moreover, we demonstrate that embigin expression can be induced by the pro-inflammatory cytokines IL-1b and Il-17. Collectively, these findings provide insight into the origin of the myofibroblasts that drive interstitial cardiac fibrosis, confirming that this population arise from cytokine stimulated resident fibroblasts, rather than an alternate non-fibroblast lineage.

We believe that our findings could have important practical and therapeutic implications. Currently, detecting and quantifying fibrosis largely relies on histological analysis, which can be insensitive (particularly in developing fibrosis), laborious and difficult to quantitate. Consequently, our identification of endogenous myofibroblast-specific markers could advance the detection myofibroblasts. Specifically, the capacity to identify myofibroblasts with high-throughput quantitative methods such as flow cytometry would greatly improve the ability to measure when myofibroblasts develops with great sensitivity and precisely quantitate this population. This capacity would be highly useful when evaluating disease progression and the efficacy of anti-fibrotic therapeutics. Moreover, the discovery of endogenous surface markers enables the isolation of live myofibroblasts (as we have done here) for further analysis. Consequently, we propose that our discovery of myofibroblast-specific markers could greatly aid future research in this field.

Our findings have potential therapeutic implications. Specifically, our demonstration that intravenously injected anti-embigin antibodies selectively bind myofibroblasts paves the way for antibody-based therapies. While preliminary studies showed that antibody opsonisation was insufficient to deplete myofibroblasts (data not shown), we propose that alternate antibody targeting therapies could be explored. For example, bi-specific T cell engaging (BiTES) antibodies targeting embigin could harness T cell killing to clear myofibroblasts, while antibody-drug conjugates could deliver cytotoxic compounds to embigin+ myofibroblasts. Alternatively, hypersialylated anti-embigin antibodies could be employed to deposit immunosuppressive antibodies within fibrotic and/or inflammatory zones. Thus, our findings provide the basis for novel strategies to target activated myofibroblasts.

Finally, we made the intriguing observation that fibroblasts in acutely injured skeletal muscle transiently upregulate embigin, but not ChL-1. These findings reveal two key points. First, our findings that embigin is expressed by cardiac and skeletal muscle fibroblasts in chronic inflammatory and acute injury suggest that embigin is a conserved marker for acutely activated fibroblasts. Second, based upon its differential expression in acute and chronic injury, we speculate that ChL-1 may only be expressed upon the terminally differentiated myofibroblasts. Hence, we believe that the combination of embigin and ChL-1 may enable the identification of intermediate (ChL-1-/emb+) and fully differentiated (ChL-1+/Emb+) myofibroblasts. However, these are preliminary observations, and further studies are required to understand the expression sequence and their implications in determining fibrotic outcomes.

In summary, our study has demonstrated that ChL-1 and embigin are upregulated by the activated myofibroblasts that drive fibrosis. Consequently, we propose that ChL-1 and embigin are novel myofibroblast-specific markers that could be used to identify, enumerate and target this pathogenic population.

## Acknowledgement

We thank Edan Azzopardi, Tom Kitson and Lauren Wilkins (WEHI, Melbourne, Australia) for outstanding technical assistance and Drs Christine Biben and Philippe Bouillet (WEHI, Melbourne, Australia) for the provision of mice.

## Sources of Funding

This work was supported by generous support from Jenny Tathcell and the Reid Charitable Trusts and grants from the Arthritis Rheumatology Australia and the Australian National Health and Medical Research Council (NHMRC; Program Grant 1113577 and Clinical Practitioner Fellowship 1154235 to I.P.W.).

## Conflict of Interest

The authors declare no conflict of interest relating to this research.

## Figure Legends

**Supplementary Table 1.**
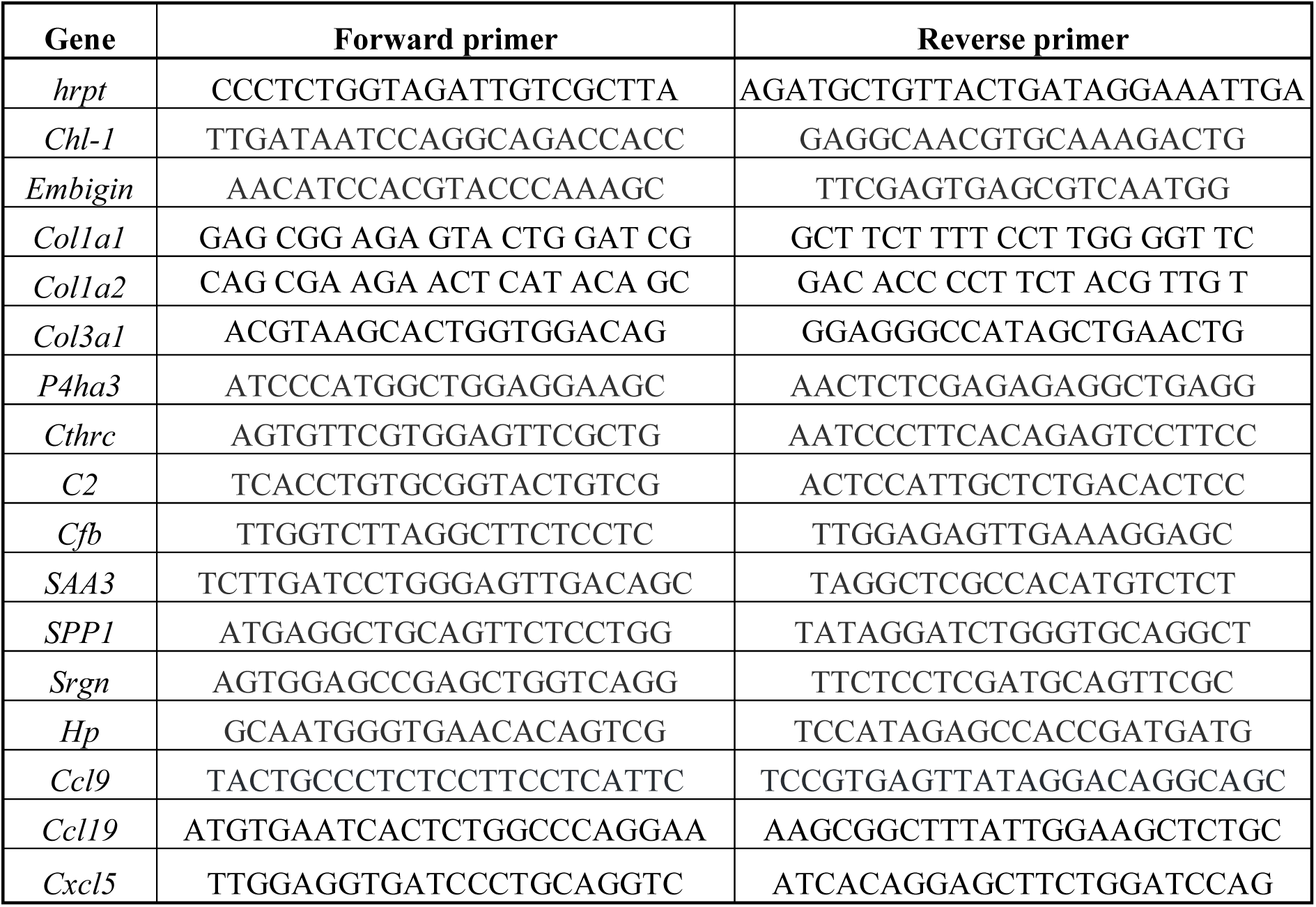
Primer sequences.

## Notes

### Competing Interest Statement

The authors have declared no competing interest.

